# DNA density is a better indicator of a nuclear bleb than lamin B loss

**DOI:** 10.1101/2024.02.06.579152

**Authors:** Samantha Bunner, Kelsey Prince, Karan Srikrishna, Emily Marie Pujadas, Antonia Amonu McCarthy, Anna Kuklinski, Olivia Jackson, Pedro Pellegrino, Shrushti Jagtap, Imuetiyan Eweka, Colman Lawlor, Emma Eastin, Griffin Yas, Julianna Aiello, Nathan LaPointe, Isabelle Schramm von Blucher, Jillian Hardy, Jason Chen, Vadim Backman, Anne Janssen, Mary Packard, Katherine Dorfman, Luay Almassalha, Michael Seifu Bahiru, A. D. Stephens

## Abstract

Nuclear blebs are herniations of the nucleus that occur in diseased nuclei that cause nuclear rupture leading to cellular dysfunction. Chromatin and lamins are two of the major structural components of the nucleus that maintain its shape and function, but their relative roles in nuclear blebbing remain elusive. Lamin B is reported to be lost in blebs by qualitative data while quantitative studies reveal a spectrum of lamin B levels in nuclear blebs dependent on perturbation and cell type. Chromatin has been reported to be decreased or de-compacted in nuclear blebs, but again the data are not conclusive. To determine the composition of nuclear blebs, we compared the immunofluorescence intensity of lamin B and DNA in the main nucleus body and nuclear bleb across cell types and perturbations. Lamin B nuclear bleb levels varied drastically across MEF wild type and chromatin or lamins perturbations, HCT116 lamin B1-GFP imaging, and human disease model cells of progeria and prostate cancer. However, DNA concentration was consistently decreased to about half that of the main nucleus body across all measured conditions. Using Partial Wave Spectroscopic (PWS) microscopy to measure chromatin density in the nuclear bleb vs body we find similar results that DNA is consistently less dense in nuclear blebs. Thus, our data spanning many different cell types and perturbations supports that decreased DNA is a better marker of a nuclear bleb than lamin B levels that vary widely.

## Introduction

Nuclear blebs are a hallmark of many human afflictions including aging, progeria, muscular dystrophy, neurological defects, and a subset of cancers (Stephens *et al*., 2019b; Kalukula *et al*., 2022). An example is prostate cancer where the levels of nuclear blebbing increase with pathology grade measured as Gleason Score (Helfand *et al*., 2012). Studies over the last 20 years have deduced the nuclear components that are essential for maintaining normal ellipsoidal nuclear shape are chromatin (Furusawa *et al*., 2015; Schreiner *et al*., 2015; Stephens *et al*., 2018, 2019a), lamins (Lammerding *et al*., 2006; Chen *et al*., 2018; Vahabikashi *et al*., 2022), and linkers within and between them (Liu *et al*., 2021; Strom *et al*., 2021; Jung-Garcia *et al*., 2023). Upon perturbation of these components the nucleus becomes weaker and is deformed by external forces from actin contraction (Nmezi *et al*., 2019; Jung-Garcia *et al*., 2023; Pho *et al*., 2023), actin compression (Le Berre *et al*., 2012; Hatch and Hetzer, 2016), transcription activity (Berg *et al*., 2023), and/or migration through constrictions (Denais *et al*., 2016; Raab *et al*., 2016; Pfeifer *et al*., 2018). This deformation leads to a herniation called a nuclear bleb. The high curvature of this bleb leads to nuclear rupture that then causes nuclear dysfunction via disruption of DNA damage/repair, transcription, and cell cycle control (Xia *et al*., 2018; Stephens, 2020; Kalukula *et al*., 2022). However, the composition of a nuclear bleb remains highly debated and elusive. A clear determination of nuclear bleb composition will provide insights into how a nuclear bleb forms and how to accurately detect it.

The nuclear lamina is composed of four separate but interacting peripheral meshworks of lamin intermediate filaments A, C, B1, and B2 (Shimi *et al*., 2015; Turgay *et al*., 2017). It has been widely reported that nuclear blebs are devoid of both B-type lamins via qualitative data, while quantitative data disagree. Some publications go as far as using loss of lamin B1 to determine what is and is not a nuclear bleb by renaming blebs lamin B less domains (LBLDs, (Helfand *et al*., 2012; Bercht Pfleghaar *et al*., 2015). Many publications provide qualitative data via one to three example images to conclude that nuclear blebs are devoid of lamin B (Yang *et al*., 2005; Shimi *et al*., 2008; Wren *et al*., 2012; Bercht Pfleghaar *et al*., 2015; Cho *et al*., 2018; Patteson *et al*., 2019; Gauthier and Comaills, 2021; Kamikawa *et al*., 2021). Recent publications have begun to quantify lamin B1 levels in nuclear blebs revealing that 25-50% of nuclear blebs are actually positive for lamin B (Stephens *et al*., 2018; Nmezi *et al*., 2019; Jung-Garcia *et al*., 2023). Thus, statements made using qualitative single images are not supported by quantitative measurements of lamin B1 levels in nuclear blebs. Instead, recent quantitative studies point to a more complicated picture of lamin B1 levels which can vary widely in nuclear blebs. We hypothesize that the disconnected nature of these studies, each working on one cell type and perturbation, leads to an incomplete view of nuclear bleb composition. Thus, nuclear bleb composition needs to be quantitatively analyzed across perturbations, cell types, live cell, and disease models.

Decreased DNA concentration in the nuclear bleb provides an alternative indictor of nuclear blebs compared to lamin B. Opposite to the mixed reporting of lamin B, nuclear blebs are more consistently reported to have less DNA in the bleb. Furthermore, enrichment of euchromatic markers has been qualitatively and quantitatively reported in nuclear blebs (Shimi *et al*., 2008; Stephens *et al*., 2018). While decreased DNA and histone modification decompaction are reported more consistently in nuclear blebs, a rigorous study across multiple perturbations and cell lines is required to determine the consistency of nuclear bleb labeling compared to lamin B1.

To determine the composition of nuclear blebs, we used immunofluorescence to label DNA and lamin B1 or B2 in mouse and human cells. Mouse embryonic fibroblasts (MEF) provide the ability to compare the composition of wild type low-percentage nuclear blebbing to chromatin or lamin perturbations previously reported to increase nuclear blebbing (Stephens *et al*., 2019b; Kalukula *et al*., 2022). In MEFs, DNA concentration displayed a consistent decrease in the nuclear bleb across wild type and perturbations while lamin B presence in the bleb was altered significantly dependent on condition. Live cell imaging of nuclear blebs and nuclear bleb formation using a CRISPR labeled GFP-lamin B1 revealed inconsistent lamin B presence in bleb while SiR-DNA concentration was consistently decreased. Next, we compared human disease cell models of progeria and prostate cancer which revealed yet again decreased DNA concentration in the nuclear bleb while lamin B1 was absent in progeria blebs and present in prostate cancer nuclear blebs. Finally, we utilized live-cell Partial Wave Spectroscopic (PWS) microscopy to confirm our results in HCT116 cells modified with the Auxin-inducible Degron System (AID) to induce simultaneous lamin B1 and lamin B2 degradation. Our results indicate that DNA density is a more reliable marker for nuclear blebs than lamin B across multiple perturbations and cell types.

## Results

### Nuclear blebs have consistently decreased DNA concentration while lamin B1 levels are highly variable depending on condition

To compare the composition of nuclear blebs and their consistency between wild type cells and those with perturbations that induce increased nuclear blebbing, we used mouse embryonic fibroblasts (MEFs). Nuclear blebs were determined by > 1 µm protrusion of the nucleus as previously defined (Stephens *et al*., 2018, 2019a; Berg *et al*., 2023; Pho *et al*., 2023) To determine the composition of nuclear blebs, we applied immunofluorescence imaging to label nuclei for DNA (DAPI), lamin B1 or B2, and nuclear pore complexes (NPCs). We measured the relative levels of each component within the nuclear bleb and compared it to the main nuclear area, referred to as the nuclear body. In wild type MEF cells the mean intensity measurements revealed significantly decreased DNA concentration in the nuclear bleb relative to body 0.55 ± 0.03 **Figure 1, A and B**). Lamin B1 staining also revealed a significant decrease in levels in the nuclear bleb compared to the body 0.62 ± 0.06. However, ratios of fluorescent signals in the nuclear bleb to body were about two-fold more variable than DNA (standard deviation DNA 0.17 vs lamin B1 0.32; **Figure 1A, C, D**). This was largely due to a small population of nuclear blebs showing no loss of lamin B1, in agreement with previous reports (Stephens *et al*., 2018). Overall, wild type nuclear blebs present an overall loss of both DNA and lamin B1 in nuclear blebs.

**Figure 1.**
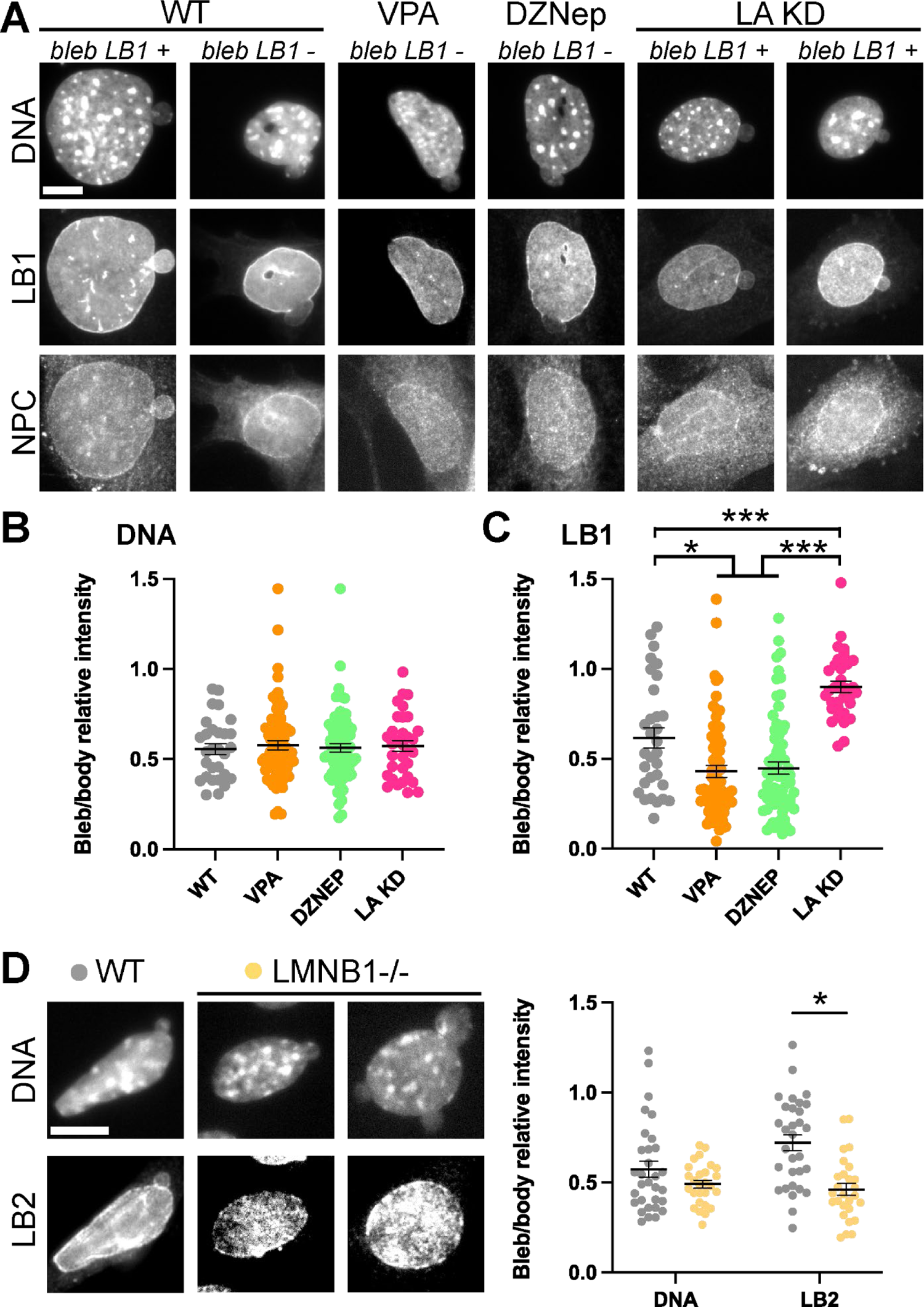
The composition of MEF wild type and perturbation-based nuclear blebs shows decreased DNA concentration but an inconsistent measure of lamin B levels. (A) Example images of mouse embryonic fibroblast (MEF) blebbed nuclei from wild type (WT), chromatin decompaction overnight drug treatments to increase euchromatin via valproic acid (VPA) or decrease heterochromatin via (DZNep), and lamin A knockdown (LA KD). Nuclei were labeled with DAPI to stain DNA, anti-lamin B1 immunofluorescence (LB1), and anti-nuclear pore complex immunofluorescence (NPC). (B, C) Graphs of bleb/body relative intensity ratio for (B) DNA and (C) LB1 for WT n = 31, VPA, n = 67, DZNep n = 71, and LA KD n = 33. (D) Example images a graph of blebbed nuclei in MEF WT (n = 32) and lamin B1 knockout (LMNB1-/-, n = 26) labeled with DAPI to stain DNA and anti-lamin B2 immunofluorescence (LB2). Graph shows bleb/body relative intensity ratio for (B) DNA and (C) LB1. Student’s t-test p values reported as *<0.05, **<0.01, ***<0.001, no asterisk denotes no significance, p>0.05. Error bars represent standard error. Scale bar = 10 µm.

Chromatin decompaction perturbations cause increased nuclear blebbing without changes in nuclear lamins (Stephens *et al*., 2018). Applying these perturbations provides an opportunity to determine if non-lamin-based nuclear perturbation causes any changes in bleb composition relative to wild type cells. Specifically, we used a histone deacetylase inhibitor valproic acid (VPA) which increases euchromatin and a histone methyltransferase inhibitor DZNep to decrease levels of heterochromatin. Both treatments that cause chromatin decompaction-induced nuclear blebs showed consistently decreased levels of DNA in the nuclear bleb relative to the body, similar to that of nuclear blebs in wild type (**Figure 1, A and B**). On the other hand, chromatin decompaction-induced nuclear blebs showed a significant decrease in lamin B1 staining relative to wild type nuclear blebs (WT 0.62 vs. VPA and DZNep 0.43 and 0.45 ± 0.03, **Figure 1, A and C**). Thus, chromatin decompaction results in similar decreased levels of DNA in the bleb compared to body as wild type, but a greater decrease in lamin B1 levels in the bleb compared to the body.

To further characterize the composition of these nuclear blebs and determine the most consistent marker for them, we repeated these experiments on nuclear blebs induced by lamin perturbation. Lamin A is usually present in nuclear blebs after nuclear rupture as it is then recruited but is absent in pre-ruptured blebs (Shimi *et al*., 2008, 2015; Denais *et al*., 2016; Kamikawa *et al*., 2021; Sears and Roux, 2022). Loss of lamin A only via constitutive shRNAi knockdown increases nuclear blebbing (Vahabikashi *et al*., 2022; Berg *et al*., 2023), and provides the ability to determine if lamin A presence or absence affects the composition of the nuclear bleb. Lamin A knockdown nuclear blebs showed a similar significant loss of DNA signal in the bleb compared to the body as in wild type cells (**Figure 1, A and B**). Interestingly, lamin A knockdown nuclear blebs showed a drastic increase in lamin B1 bleb/body ratio compared to wild type nuclear blebs (WT 0.62 vs. LA KD 0.90 ± 0.03, p < 0.001, **Figure 1, A and B)**. Thus, lamin A knockdown blebs showed on average no significant loss of lamin B1 in the nuclear bleb relative to the nuclear body (p > 0.05). Taken together, nuclear blebs in wild type, chromatin perturbations, and lamin A knockdown have similar levels of decreased DNA in the bleb while each condition has a significantly different level of lamin B1 in nuclear blebs.

### Nuclear pore complex presence or absence in the nuclear bleb varied based on lamins

In a continued effort to understand the composition of nuclear blebs, we also imaged the nuclear pore complex in the bleb compared to the body. As NPCs ensure macromolecular transport between the nucleus and cytoplasm (Beck and Hurt, 2017), their presence or absence in nuclear blebs could provide insights into the functionality of nuclear blebs. Interestingly, both wild type cells and cells treated with VPA or DZNep to induce chromatin decompaction showed a qualitative correlation between lamin B1 loss and NPC presence in nuclear blebs (**Figure 1A**). Oppositely, lamin A knockdown nuclear blebs contain lamin B1, but are devoid of NPCs. This data suggests that NPC presence in nuclear blebs is highly dependent on both lamin A and lamin B. Visually, we did not detect a clear trend in NPC staining as a marker of nuclear blebs in comparison to DNA staining, which further suggests that DNA density is a more reliable marker for these nuclear herniations.

### Lamin B2 levels are similarly heterogenous in nuclear blebs and can be altered by perturbations while DNA concentration in nuclear bleb remains consistent

Lamin B1 loss was one of the first reported perturbations that results in increased nuclear blebbing (Lammerding *et al*., 2006; Vargas *et al*., 2012; Shimi *et al*., 2015; Hatch and Hetzer, 2016). To determine if lamin B1 loss alters the composition of the nuclear blebs, we applied immunofluorescence imaging to label DNA and lamin B2 in both wild type and lamin B1 knockout (LMNB1-/-) MEF cells. Again, DNA concentration was consistently decreased in nuclear blebs relative to the body in wild type cells (0.57 ± 0.05, **Figure 1D**). As was the case for other nuclear perturbations that increased nuclear blebbing, lamin B1 knockout cells also displayed a relatively consistent decrease in DNA in the nuclear bleb relative to the nuclear body (0.49 ± 0.02, **Figure 1D**). Thus, decreased DNA concentration in nuclear blebs were measured consistently for wild type and all chromatin and lamin perturbations.

Next, we looked at lamin B2 presence in nuclear blebs. Like lamin B1, lamin B2 levels in nuclear blebs also varied in wild type cells, with some lower and some similar to that of the nuclear body. In lamin B1 knockout cell, lamin B2 displayed a significant decrease in nuclear blebs relative to wild type (WT 0.71 ± 0.05 vs. LMNB1-/- 0.47 ± 0.03, p < 0.001, **Figure 1D**). This data suggests that while nuclear blebs show a clear and significant loss of DNA across perturbations, the levels of lamin B2 are dependent on lamin B1 (**Figure 1D**) which we show is dependent on both histone modification state and lamin A levels (**Figure 1A-C**).

### Live cell imaging recapitulates the consistent loss of DNA and variable lamin B1 presence in nuclear blebs

To further explore nuclear bleb composition, we used HCT116 colon cancer cells with a CRISPR labeled endogenous GFP-lamin B1. First, we performed immunofluorescence staining for both DNA (DAPI) and lamin B2 to compare with GFP-lamin B1. Similar to MEFs we investigated rare nuclear blebbing in untreated cells and increased nuclear blebbing via VPA-treatment to decompact chromatin. Immunofluorescence imaging recapitulated major findings form MEFs as DNA concentration was consistently decreased in nuclear blebs while both lamin B1 and B2 levels varied drastically (**Figure 2A**). To verify that fixed-cell immunofluorescence does not cause artifacts, we next imaged nuclear bleb composition in live cells. Live cell imaging of GFP-lamin B1 and Hoechst-stained DNA revealed that both wild type untreated and VPA-treated nuclear blebs present a consistent decreased level of DNA while lamin B1 level varied widely (**Figure 2B**), a recapitulation of the data obtained via fixed-cell immunofluorescence. Both MEF and human HCT116 nuclear blebs show a clear and consistent loss of DNA while lamin B levels vary.

**Figure 2.**
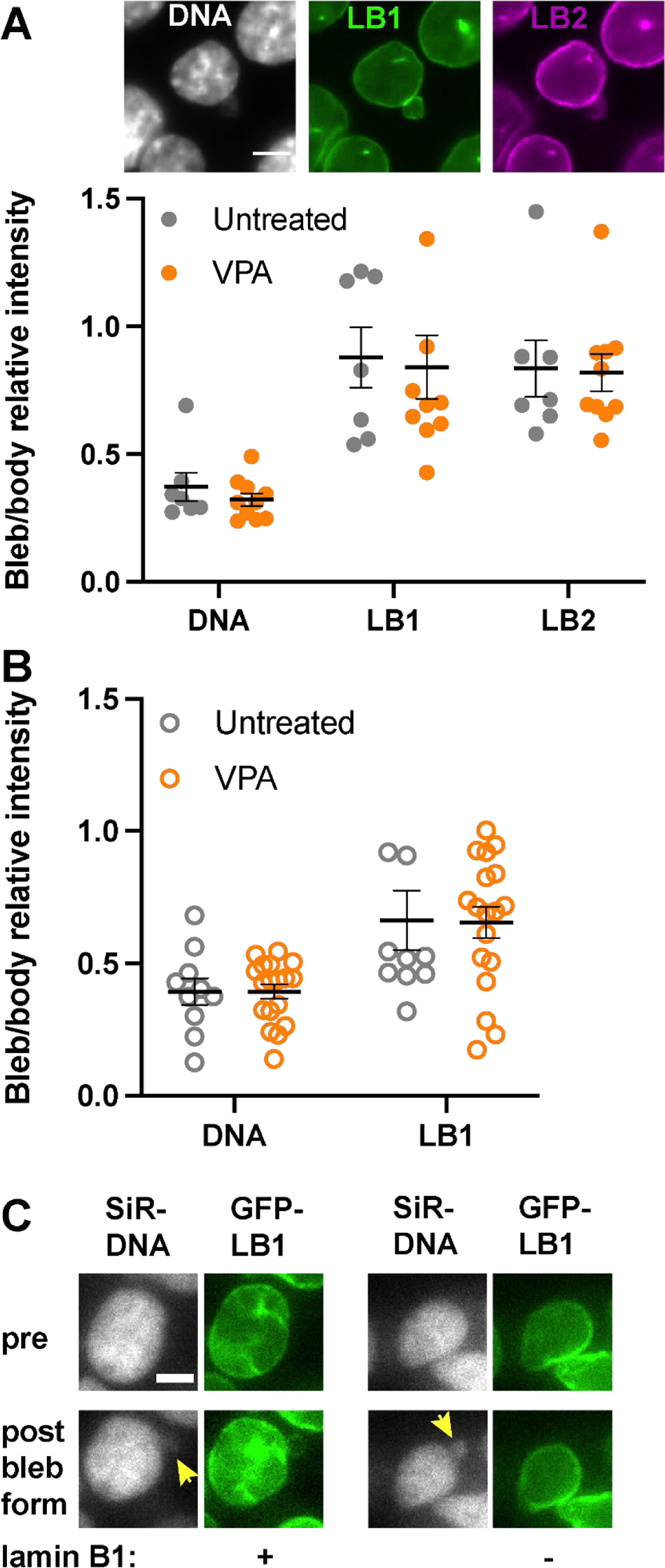
Immunofluorescence and live cell imaging of HCT116 cells with GFP-lamin B1 reproduces variable lamin B is presence nuclear blebs while DNA remains consistently decreased. (A) Example image and graph of relative levels in the nuclear bleb determined by immunofluorescence imaging of HCT116 cells with CRISPR labeled GFP-lamin B1 with Hoechst labeled DNA and anti-lamin B2 labeling in untreated (n = 7) and VPA-treated (n = 10). (B) Graph of live cell imaging levels of Hoechst-stained DNA and GFP-lamin B1 in the nuclear bleb relative to the nuclear body in untreated (n = 13) and VPA-treated (n = 18). (C) Example images from live cell timelapse imaging of nuclear bleb formation in HCT116 GFP-lamin B1 cells with DNA stained by SiR-DNA shows nuclear blebs can form either with or without lamin B1. Error bars represent standard error. Scale bar = 5 µm.

Studies have postulated that localized loss of lamin B1 aids nuclear bleb formation(Lammerding *et al*., 2006). Along these lines, lamin B1 loss could be temporary during nuclear bleb formation which might lead to newly formed nuclear blebs being absent for lamin B1 while long-lived nuclear blebs have lamin B1. To test the hypothesis, we imaged nuclear bleb formation in live HCT116 GFP-lamin B1 nuclei stained with SiR-DNA. We find that nuclear blebs can form with or without the presence of lamin B1, suggesting the variance of lamin B1 levels in nuclear blebs occurs during formation. (**Figure 2C**). Even in live cell imaging of nuclear bleb formation, SiR-DNA is decreased in the nuclear bleb upon formation. Taken together, both live cell and static immunofluorescence images of nuclear blebs as well as live cell imaging of bleb formation show a consistent loss of DNA and variable lamin B1 presence.

### Different cell types and models of human disease have a similar loss of DNA in the nuclear bleb, while lamin B1 levels change drastically

Based on the consistent results we obtained in MEFs and HCT116 cells, we next tested the reliability of decreased DNA as a marker for blebbing in different disease models that are associated with increased nuclear deformation and blebbing: human progeria disease and prostate cancer cell lines. Progeria Syndromes are rare genetic disorders which recapitulate many aspects of normal aging and are mainly caused by mutations in DNA repair proteins or proteins associated with the nuclear envelope (Butin-Israeli *et al*., 2012). Further, loss of lamin B1 in nuclear blebs has frequently been reported in progeria disease model cell lines (Yang *et al*., 2005; Shimi *et al*., 2008; Bercht Pfleghaar *et al*., 2015; Cho *et al*., 2018; Gauthier and Comaills, 2021). The use of prostate cancer cell line models also provides the ability to assay lamin B1 presence in a different model of human disease.

To determine quantitative levels of lamin B1 in Progeria-based nuclear blebs, we used wild type (hTERT) and and Nestor-Guillermo progeria syndrome (NGPS) immortalized fibroblasts to measure DAPI and lamin B1 relative bleb levels. NGPS is caused by a homozygous A12T point mutation in Barrier-to-autointegration-Factor1 (BAF), a small protein binding to DNA, A-type lamins and several nuclear envelope proteins(Cabanillas *et al*., 2011; Puente *et al*., 2011). The disease mutation decreases the interaction between A-type lamins and BAF (Janssen *et al*., 2022a) resulting in a lack of A-type lamin recruitment to nuclear envelope ruptures. NGPS patient cell blebs were thus reported to lack A-type lamins (Janssen *et al*., 2022a). Analysis of DNA concentration showed significantly decreased DNA labeling intensity in the nuclear bleb relative to the nuclear body consistently while not changing between control (hTERT) and progeria (NGPS) human fibroblasts (**Figure 3, A and B**) or prostate cancer cell lines **(Figure 3, A and C)**. Oppositely, Lamin B1 levels varied drastically between the two different model cell lines (**Figure 3, D and E**). Lamin B1 was consistently absent in nuclear blebs compared to the nuclear body in both wild type and NGPS fibroblasts (**Figure 3, A and D**), suggesting lamin B1 loss in nuclear blebs is not specifically due to the BAF mutation. In most prostate cancer cells lamin B1 was present in nuclear blebs at similar levels to the rest of the nucleus (**Figure 3E**). However, a minor subpopulation of cells did show near complete loss in some blebs. Interestingly, different stages of prostate cancer from the less aggressive LNCaP to more aggressive DU145 showed increased nuclear blebbing and loss of nuclear shape but maintained similar levels of lamin B1 in the bleb. These results confirm not only that loss of DNA concentration is a reliable indicator across different cell types, but also that between cell types lamin B1 levels in nuclear blebs vary drastically.

**Figure 3.**
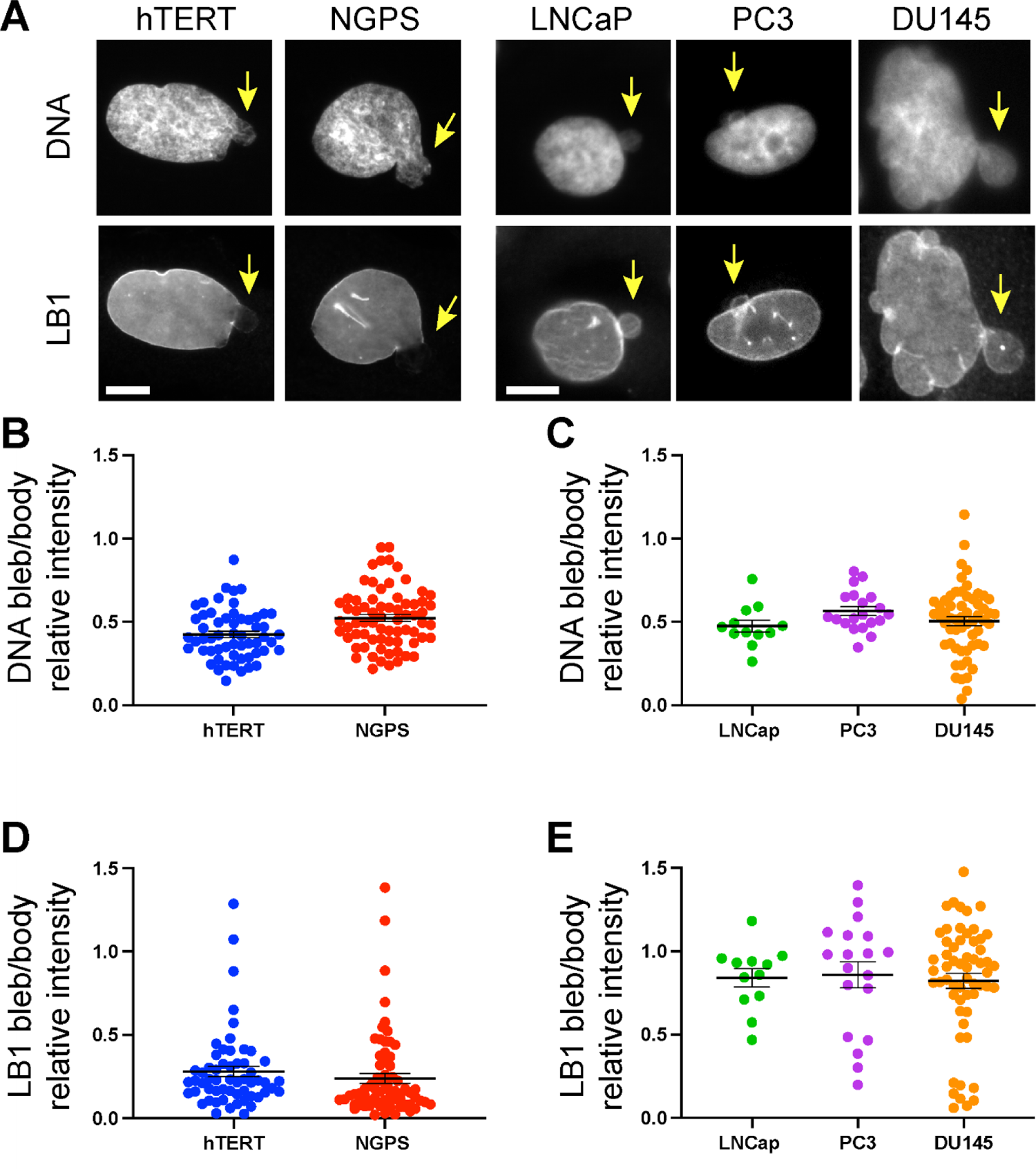
Progeria and prostate cancer model cell lines recapitulate decreased DNA concentration in nuclear blebs while lamin B1 is drastically different. (A) Example images of a blebbed nucleus in hTERT and NGPS human fibroblasts along with human prostate cancer model cell lines LNCaP, PC3, and DU145. Nuclei are labeled with DAPI to stain DNA and anti-lamin B1 immunofluorescence (LB1). The yellow arrow denotes nuclear bleb. Graphs of bleb/body relative intensity ratio for (B) DNA and (C) LB1 for human wild type (hTERT, n = 58) and Progeria (NGPS, n = 71) fibroblast. Graphs of bleb/body relative intensity ratio for (D) DNA and (E) LB1 human prostate cancer cells of increasing aggressiveness LNCaP n = 12, PC3 n = 20, and DU145 n = 33. Student’s t-test p values reported as *<0.05, **<0.01, ***<0.001, no asterisk denotes no significance, p>0.05. Error bars represent standard error. Scale bar = 10 µm.

### Loss of DNA density is a consistent metric of nuclear blebbing across perturbations

Finally, we assessed if the results obtained with immunofluorescence imaging would still hold using a more high-throughput technique that can detect statistical changes in chromatin structure across entire cell populations. Partial Wave Spectroscopic (PWS) microscopy provides label-free measurements of nanoscale structural changes with a sensitivity to structures between ∼ 20 and 200 nm in live cells without the use of exogenous labels (Almassalha *et al*., 2016; Gladstein *et al*., 2018, 2019). To do this, PWS directly measures variations in spectral light interference that results from light scattering. As this scattering is due to heterogeneities in chromatin density (Li *et al*., 2021), PWS provides a way to confirm that reduced DNA density is a reliable marker for nuclear blebbing. Chromatin packing scaling (D) can be calculated from these fluctuations in chromatin density. To obtain D measurements within nuclear blebs and nuclear bodies, we used HCT116 cells modified with the AID system to induce simultaneous rapid lamin B1 and lamin B2 degradation (Nishimura *et al*., 2009; Yesbolatova *et al*., 2019; Pujadas *et al*., 2023), termed _HCT116_LMN(B1&B2)-AID cells.

Using PWS, we confirmed that HCT116^LMN(B1&B2)-AID^ cell nuclei showed a significant decrease in D in the nuclear bleb relative to the nuclear body (**Figure 4**), similar to the observed decrease in Hoechst labeled fluorescent measures seen in HCT116 (**Figure 2**). This decrease in D was shown for both control and auxin-induced simultaneous degradation of lamin B1 and lamin B2 (**Figure 4A**). To confirm these results in cells with decompacted chromatin, we then treated HCT116^LMN(B1&B2)-AID^ cells with GSK343, a potent EZH2 inhibitor that specifically prevents H3K27 methylation, and Trichostatin A (TSA), a specific HDAC class I and II inhibitor. Our results demonstrated significant decreases in D within nuclear blebs of DMSO-treated control, GSK343-treated, and TSA-treated cells in comparison to nuclear main nuclear body (**Figure 4B**). These differences were apparent in representative D maps of each condition (**Figure 4C-D**). These results demonstrate that in live-cells, either with or without the presence of B-type lamins, with EZH2 inhibition, and with HDAC inhibition, reduced DNA or chromatin density is a consistent marker of nuclear blebbing. Taken together, this work establishes a more reliable method of quantifying and characterizing nuclear blebs across multiple perturbations and cell types. We therefore provide a more complete assessment of nuclear bleb composition.

**Figure 4.**
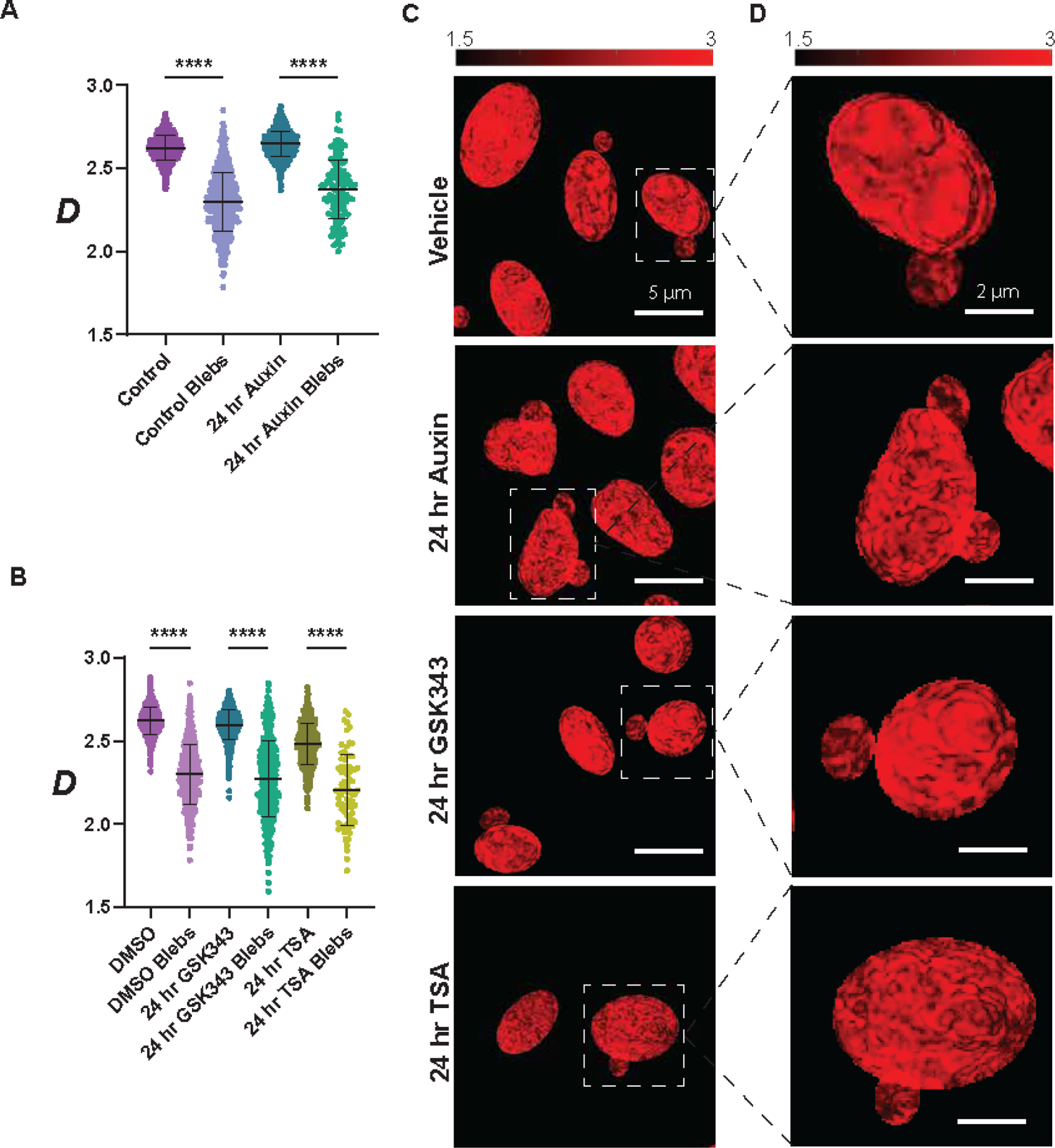
Partial wave spectroscopy reveals a consistent loss of DNA density in nuclear blebs. **A.** Live-cell PWS microscopy demonstrates that average D is much lower in the nuclear bleb than the nuclear body in untreated (Control) and auxin-treated HCT116^LMN(B1&B2)-AID^ cells (N > 100 cells/ condition for each cell type; *P ≤ 0.05, **P ≤ 0.01, ***P ≤ 0.001, ****P ≤ 0.0001) **B.** Live-cell PWS microscopy demonstrates that average D is much lower in the nuclear bleb than the nuclear body upon GSK343 or TSA treatment compared to the 24 hr DMSO treatment (vehicle control). Violin plots show the median and quartiles for the unpaired two-tailed t-test between selected groups. (N > 100 cells/ condition for each cell type; *P ≤ 0.05, **P ≤ 0.01, ***P ≤ 0.001, ****P ≤ 0.0001). **C.** Representative PWS D maps of several cells with nuclei **D.** and zoomed-in images for each cell type are shown. Scale bars represent 5 µm and 2 µm, respectively.

## Discussion

Nuclear blebs are both hallmarks and a major cause of human diseases. Even though their importance is clear, we still understand very little about nuclear blebs. A striking example is the lack of understanding around nuclear bleb composition and a consistent marker for nuclear blebs. Previous, qualitative reports focused on a few images showing a loss of lamin B in nuclear blebs. However, data standards now require quantification which revealed that lamin B1 can vary widely. In this report, we used fluorescence imaging to quantify the levels of both DNA and lamin B1 in nuclear blebs across four different cell types and many more perturbations. We provide quantitative data showing how drastically lamin B1 levels can differ in nuclear blebs due to either cell type or perturbation, among other possible factors. On the other hand, we find that, relative to the main nucleus, nuclear blebs consistently display decreased DNA concentration across cell types, perturbations, and levels of disease progression. Our analysis reveals that these findings are consistent between different imaging modalities by using PWS to measure the statistical chromatin density. Our work provides a change in the paradigm of how nuclear blebs might be quantitatively identified by DNA density and not by lamin B1 levels.

### Lamin B levels in nuclear blebs are highly dependent on cell type and perturbation

Previous studies of nuclear bleb composition in progeria were pivotal in building our current understanding of this important phenomena that is both a hallmark and cause of human disease. Several groups reported that across different progeria types, lamin B1 and B2 were absent in nuclear blebs based on qualitative data (Shimi *et al*., 2008; Butin-Israeli *et al*., 2012; Bercht Pfleghaar *et al*., 2015). Our data suggests that lamin B loss reported in previous papers focused on progeria models is likely due to the use of human fibroblasts which quantitatively show little to no lamin B in nuclear blebs (**Figure 3**). It is important to note that previous qualitative data are supported by quantitative data, but that the generalization of lamin B loss to other cell types and/or perturbations in blebs is not. This data is heavily contrasted by mouse fibroblasts, human colon cancer cells (HCT116), and many prostate cancer cells (LNCaP, PC3, DU145) which display varied or unchanged levels of lamin B in nuclear blebs (**Figures 1-3**). Furthermore, we reveal that lamin B1 levels in the nuclear bleb can be altered based on chromatin or lamin perturbation (**Figure 1**). Thus, in agreement with other quantitative studies of nuclear bleb composition (Stephens *et al*., 2018; Nmezi *et al*., 2019; Janssen *et al*., 2022a, 2022b; Jung-Garcia *et al*., 2023), lamin B1 levels can vary widely and depend on cell type and perturbation.

### The ever-growing importance of DNA/chromatin in nuclear blebbing

Chromatin is now a well-established determinant of nuclear blebs (Furusawa *et al*., 2015; Stephens *et al*., 2018, 2019a; Kalinin *et al*., 2021). Isolated single nucleus micromanipulation force measurements provide the ability to separate chromatin and lamin mechanical contribution to nuclear force resistance (Stephens *et al*., 2017; Currey *et al*., 2022) recapitulated by AFMLS (Hobson *et al*., 2020). Chromatin acts as the initial and main mechanical component of the nucleus to resist antagonistic external cytoskeletal forces (Pho *et al*., 2023) and internal transcriptional chromatin movement (Berg *et al*., 2023) that drive nuclear blebbing which leads to nuclear rupture and dysfunction (Stephens *et al*., 2019b; Stephens, 2020; Kalukula *et al*., 2022). Thus, it is not surprising that decreased DNA concentration provides a strong indicator of nuclear blebs.

Overall, decreased DNA concentration in the nuclear bleb provides the most consistent indicator to date. Decreased DNA in the nuclear bleb is consistent across four different cell types, multiple perturbations, or levels of disease progression. More specifically, decreased concentration of DNA in the nuclear blebs was demonstrated in both untreated wild type cells, which show low percentage of nuclear blebbing, and upon perturbation by chromatin decompaction (VPA, TSA, DZNep, GSK343) or lamins (LA KD, LMNB1-/-, GFP-AID-lamin B1) which display high percentage of nuclear blebbing (**Figure 1**). This suggests that decreased DNA density is a hallmark of nuclear blebbing while bleb composition is independent of both global histone modification state and lamins. Thus, this data supports the idea that DNA and/or chromatin decreased density is an essential factor driving nuclear bleb formation. At the least, nuclear bleb formation is not dependent on lamin B1 composition as example timelapse imaging shows that nuclear blebs form with or without lamin B1 (**Figure 2C**). This same data shows a decreased level of DNA in the nuclear bleb relative to the nuclear body. Finally, wild type to progeria (NGPS) fibroblasts and less aggressive (LNCap) to more aggressive (DU145) prostate cancer also has no effect on DNA concentration in nuclear blebs. Taken together, the consistency of decreased DNA concentration in nuclear blebs strongly supports chromatin decompaction as a driving mechanism in nuclear bleb formation.

### Nuclear bleb composition informs possible models of nuclear bleb formation

The composition of the nuclear bleb may aid in uncovering the steps during nuclear bleb formation, a process that remains elusive. There are a few theoretical models for nuclear bleb formation when considering very simple parameters of chromatin and lamins. Chromatin could simply push out the nuclear lamina/envelope and maintain similar levels of lamin proteins. This theoretical model has data to support it in that lamin B1 positive blebs appear to have a completely intact lamina around the main nuclear body and the nuclear bleb (**Figure 1-3**). A slight deviation of this idea is that chromatin could flow through a small hole, or holes, in the lamina resulting in pushing out some lamina but not others to provide a decreased level of lamin proteins but not complete loss. This model could represent data showing that nuclear blebs can have varying levels if lamin B1 but would require more fine-tuned experiments to support as a driving model. A third model is that the lamina breaks or has a hole in which chromatin flows through to form the nuclear bleb. Previous, reports noted that lamin B1 gaps in the nuclear lamina could allow for chromatin to flow out to form a nuclear bleb. Our data support this model as lamin B1 negative nuclear blebs appear to have a sizeable gap between the nuclear body lamina and the nuclear bleb devoid of lamin B. Taken together, our data supports that many different theoretical models of chromatin and lamin interactions can arise to form a nuclear bleb of various lamin compositions, but the only uniting mechanism is less dense DNA.

### Future directions

Nuclear bleb composition, formation, and consequences remain a valuable topic for discovery. This work has deduced the consistency of DNA to label nuclear blebs, underlying its importance likely in nuclear bleb formation. However, many questions remain surrounding histone modification states that dictate chromatin compaction. Can investigating nuclear bleb composition inform the role of euchromatin, facultative heterochromatin, and constitutive heterochromatin in nuclear bleb formation? Our findings on the variable presence of lamin B in nuclear blebs raises many questions about other nuclear lamina and envelope proteins such as lamin A and C, nuclear pore complexes (NPCs), emerin, BAF, LAP2, LBR, and many more. Through leveraging our understanding of nuclear bleb composition, it is possible to gain a deeper understanding of nuclear bleb formation, a prominent even in human disease.

## Materials and Methods

### Cell Culture

Mouse Embryonic Fibroblast (MEF) were previously described in (Shimi *et al*., 2008; Stephens *et al*., 2018; Vahabikashi *et al*., 2022). MEF wild-type (WT), lamin A knockdown (LA KD), and lamin B1 knockout (LMNB1-/-) cells were cultured in DMEM (Corning) completed with 10% fetal bovine serum (FBS, HyClone) and 1% penicillin/streptomycin (Corning). Cells were grown and incubated at 37 C and 5 % CO2, passaged every 2 to 3 days.

Three prostate cancer cell lines were used: LNCaP, DU145, and PC3. DU145 and LNCaP cells were cultured in RPMI 1640 with 10% fetal bovine serum (FBS) and 1% penicillin/ streptomycin. PC3 cells were cultured in DMEM completed with 10% fetal bovine serum (FBS) and 1% penicillin. HCT116^LMN(B1&B2)-AID^ cells were grown in McCoy’s 5A Modified Medium (#16600-082, Thermo Fisher Scientific, Waltham, MA) supplemented with 10% FBS (#16000-044, Thermo Fisher Scientific, Waltham, MA) and penicillin-streptomycin (100 μg/ml; #15140-122, Thermo Fisher Scientific, Waltham, MA). To create these cells, HCT116 cells (ATCC, #CCL-247) were tagged with the AID system as previously described (Pujadas *et al*., 2023). All cells were cultured under recommended conditions at 37°C and 5% CO2. All cells in this study were maintained between passage 5 and 20. Cells were allowed at least 24 h to re-adhere and recover from trypsin-induced detachment. All imaging was performed when the surface confluence of the dish was between 40–70%. All cells were tested for mycoplasma contamination (ATCC, #30-1012K) before starting perturbation experiments, and they have given negative results.

NGPS cells (NGPS5787) and control fibroblasts (AG10803) were immortalized with SV40LT and TERT. These immortalized cell lines were a gift from Carlos López-Otín and were cultured in DMEM containing 10% fetal bovine serum and penicillin/streptomycin. Cells were maintained at 37°C and 5 % CO2.

### Biochemical Treatments

MEF WT cells were treated with either 4 mM valproic acid (VPA, 1069-66-5, Sigma) or 1µM 3-deazaneplanocin (DZNep, (Miranda *et al*., 2009)) for 24 hours before fixation and immunofluorescence.

HCT116^LMN(B1&B2)-AID^ cells were plated at 50,000 cells per well of a 6-well plate (Cellvis, P12-1.5H-N). To induce expression of OsTIR1, 2 μg/ml of doxycycline (Fisher Scientific, #10592-13-9) was added to cells 24 hours prior to auxin treatment. 1000 μM Indole-3-acetic acid sodium salt (IAA, Sigma Aldrich, #6505-45-9) was solubilized in RNase-free water (Fisher Scientific, #10-977-015) before each treatment as a fresh solution and added to HCT116^LMN(B1&B2)-AID^ cells.

HCT116^LMN(B1&B2)-AID^ cells were plated at 50,000 cells per well of a 6-well plate (Cellvis, P12-1.5H-N). Cells were given at least 24 hours to re-adhere before treatment. VPA was dissolved in media and cells were treated with 4 mM. GSK343 (Millipore Sigma, #SML0766) was dissolved in DMSO to create a 10 mM stock solution. This was further diluted in complete cell media to a final treatment concentration of 10 µM. TSA (Millipore Sigma, #T1952) was diluted in complete cell medium and added to cells at a final treatment concentration of 300 nM.

For live cell imaging, cells were treated with 1 µm sirDNA (Cytoskeleton, CY-SC007) and 1 µm verapamil (Cytoskeleton, CY-SCV01)

### Immunofluorescence

Cells were grown on coverslips in preparation. Cells were fixed with 3.2% paraformaldehyde and 0.1% glutaraldehyde in phosphate buffered saline (PBS) for 10 minutes. Between steps cells were washed with PBS Tween 20 (0.1%) and Azide (0.2 g/L) three times, denoted PBS-Tw-Az. Next, cells were permeabilized by 0.5% Triton-X 100 in PBS for 10 minutes. Again, cells were washed with PBS-Tw-Az. Humidity chambers were prepared using petri dishes, filter paper, distilled water, and parafilm. 50µL of primary antibody solution was applied to the parafilm and the cell-side and the coverslip was placed on top to incubate for 1 hour at 37C in the humidity chamber. Primary antibodies used were Mouse monoclonal Anti-Nuclear Pore Complex Proteins at 1:1,000 (Mab414, Abcam ab24609), Rabbit polyclonal Anti-Lamin B1 at 1:1,000 (Abcam ab16048), Rabbit recombinant monoclonal Anti-Lamin B2 at 1:1,000 (Abcam ab151735). The humid chambers were removed from the incubator and the coverslips were washed in PBS-Tw-Az. More humidity chambers were made for the secondary incubation. 50µL of secondary antibody was placed on the parafilm, and the coverslips were placed cell-side down. The secondary antibody solution contained goat anti-mouse antibodies FITCI at 1:1000 dilution and goat anti-rabbit dilution TRITCI at 1:1000 dilution (from Cell Signaling Technologies) and incubated in humidity chambers at 37°C for 30 minutes. After, coverslips were washed with PBS-Tw-Az. The coverslips were placed cell-side down on a slide with a drop of mounting media containing DAPI. Slides were allowed to cure for four days at 4C before imaging.

As previously described in (Janssen *et al*., 2022a, 2022b) hTERT and NGPS cells were fixed at room temperature for 10 min with 4% PFA. Cells were washed in PBS, permeabilized using 0.2% Triton-X100, and blocked using 2% bovine serum albumin (BSA) in PBS for 30 min. Cells were incubated overnight at 4◦C or for 1 h at RT in 2%BSA PBS containing primary antibody mouse anti-lamin B1 (Santa Cruz, sc-365214, 1/500). Cells were washed using PBS and incubated for 1 h at room temperature with secondary antibody in 2% BSA PBS. Cells were washed in PBS and mounted using Prolong Gold (Thermo Fischer). Images were taken on Zeiss Axio Imager Z2 using a 63× oil immersion objective (Plan APO, NA 1.4, Zeiss).

### Immunofluorescence Imaging

Immunofluorescence images were acquired using a QICAM Fast 1394 Cooled Digital Camera, 12-bit, Monochrome CCD camera (4.65 x 4.65 µm pixel size and 1.4 MP, 1392 x 1040 pixels) using Micromanager and a 40x objective lens on a Nikon TE2000 inverted widefield fluorescence microscope. Cells were imaged using transmitted light to find the optimal focus on the field of view to observe the nuclear blebs. Ultraviolet light (excitation 360 nm) was used to visualize DNA via DAPI, blue, fluorescent light (excitation 480 nm) was used to visualize nuclear pore complex (NPC) and green fluorescent light (excitation 560 nm) was used to visualize Lamin B1 or B2. Images were saved and transferred to Fiji imaging software for analysis (Schindelin *et al*., 2012).

### Nuclear Bleb Analysis

Images were exported to FIJI (Schindelin *et al*., 2012) to analyze the intensity of each component normalizing bleb intensity by nuclear body intensity. The body, bleb, and background of each nucleus image were measured by drawing regions of interest (ROI) via the polygon selection tool. Measurements of mean light intensity in each excitation light were recorded and exported to excel. Within excel, the background intensity was subtracted from the body and bleb. Then the bleb intensity was divided by the nuclear body intensity to give a measure relative to 1 for each nuclear bleb measured. These values were transferred to Prism where Mann-Whitney test, unpaired t tests, or one way ANOVA were performed, and the data was graphed.

### PWS Image Acquisition and Analysis

For live-cell measurements, cells were imaged and maintained under physiological conditions (5% CO2 and 37°C) using a stage-top incubator (In Vivo Scientific, Salem, SC; Stage Top Systems). Live-cell PWS measurements were obtained using a commercial inverted microscope (Leica, DMIRB) using a Hamamatsu Image-EM charge-coupled device (CCD) camera (C9100-13) coupled to a liquid crystal tunable filter (LCTF, CRi VariSpec) to acquire monochromatic, spectrally resolved images ranging from 500-700 nm at 2-nm intervals as previously described (Almassalha *et al*., 2016; Gladstein *et al*., 2018, 2019). Broadband illumination is provided by a broad-spectrum white light LED source (Xcite-120 LED, Excelitas). The system is equipped with a long pass filter (Semrock BLP01-405R-25) and a 63x oil immersion objective (Leica HCX PL APO). All cells were given at least 24 hours to re-adhere before treatment (for treated cells) and imaging. Briefly, PWS measures the spectral interference signal resulting from internal light scattering originating from nuclear chromatin. This is related to variations in the refractive index (RI) distribution (Σ) (extracted by calculating the standard deviation of the spectral interference at each pixel), characterized by the chromatin packing scaling (D). D was calculated using maps of Σ, as previously described (Gladstein *et al*., 2019; Eid *et al*., 2020; Virk *et al*., 2020). Measurements were normalized by the reflectance of the glass medium interface (i.e., to an independent reference measurement acquired in a region lacking cells on the dish). This allows us to obtain the interference signal directly related to RI fluctuations within the cell. Although it is a diffraction-limited imaging technique, PWS can measure chromatin density variations because the RI is proportional to the local density of macromolecules (e.g., DNA, RNA, proteins). Therefore, the standard deviation of the RI (Σ) is proportional to nanoscale density variations and can be used to characterize packing scaling behavior of chromatin domains with length scale sensitivity around 20 – 200 nm, depending on sample thickness and height. Changes in D resulting from each condition are quantified by averaging cells, taken across 3 technical replicates. Average D was calculated by first averaging D values from PWS measurements within each cell nucleus and then averaging these measurements over the entire cell population for each treatment condition.

## Acknowledgements

We would like to thank Mai Pho for technical assistance and Edward J. Banigan for helpful and insightful discussions. We would also like to thank HHMI which purchased microscopes used in Bioimaging class via a grant and The Biology Department at UMass Amherst for use of the facilities associated with ISB 360. ADS are supported by the Pathway to Independence Award (R00GM123195) and Center for 3D Structure and Physics of the Genome 4DN2 grant (1UM1HG011536). The authors declare not competing interests. This work was supported by NSF grants EFMA-1830961 and EFMA-1830969 and NIH grants R01CA228272, U54 CA268084, and U54 CA261694. Philanthropic support was generously received from Rob and Kristin Goldman, the Christina Carinato Charitable Foundation, Mark E. Holliday and Mrs. Ingeborg Schneider, and Mr. David Sachs. AJ was supported by a FEBS Long Term Fellowship. We would like to thank Carlos Lopez-Otin for providing us with the Nestor–Guillermo progeria cell lines

